# Causal associations between potentially modifiable risk factors and the Alzheimer’s disease phenome: A Mendelian randomization study

**DOI:** 10.1101/689752

**Authors:** Shea J Andrews, Brian Fulton-Howard, Paul O’Reilly, Lindsay A Farrer, Jonathan L Haines, Richard Mayeux, Adam C Naj, Margaret A Pericak-Vance, Gerard D Schellenberg, Li-San Wang, Edoardo Marcora, Alison M Goate

## Abstract

**Objective:** To evaluate the causal association of 22 previously reported risk factors for Alzheimer’s disease (AD) on the “AD phenome”: AD, AD age of onset (AAOS), hippocampal volume, cortical surface area and thickness, cerebrospinal fluid (CSF) levels of Aβ_42_, tau, and ptau_181_, and the neuropathological burden of neuritic plaques, neurofibrillary tangles, and vascular brain injury (VBI).

**Methods:** Polygenic risk scores (PRS) for the 22 risk factors were computed in 26,431 AD cases/controls and the association with AD was evaluated using logistic regression. Two-sample Mendelian randomization was used to evaluate the causal effect of risk factors on the AD phenome.

**Results:** PRS for increased education and diastolic blood pressure were associated with reduced risk for AD. PRS for increased total cholesterol and moderate-vigorous physical activity were associated with an increased risk of AD. MR indicated that only Education was causally associated with reduced risk of AD, delayed AAOS, and increased cortical surface area and thickness. Total-and LDL-cholesterol levels were causally associated with increased neuritic plaque burden, while diastolic blood pressure and pulse pressure are causally associated with increased risk of VBI. Furthermore, total cholesterol was associated with decreased hippocampal volume; smoking initiation and BMI with decreased cortical thickness; and sleep duration with increased cortical thickness.

**Interpretation:** Our comprehensive examination of the genetic evidence for the causal roles of previously reported risk factors in AD using PRS and MR, supports a causal role for education, blood pressure, cholesterol levels, smoking, and BMI with the AD phenome.

## Introduction

Late-onset Alzheimer’s disease (AD) is a debilitating neurological condition characterized by progressive deterioration in cognitive function resulting in functional decline ^1^. The primary neuropathological hallmarks of AD are the aggregation of extracellular amyloid-β (Aβ) peptides into amyloid plaques and of intracellular hyperphosphorylated tau into neurofibrillary tau tangles (NFTs), accompanied by gliosis and neurodegeneration ^1^.

In the absence of any disease-modifying therapies, the number of people living with dementia in the USA is expected to exceed 13.8 million by 2050 ^1^. Observational studies have identified potentially modifiable risk factors that could be targeted in intervention studies to reduce the risk of dementia or delay its onset, thereby significantly reducing the population prevalence of AD and related dementias ^2^. From these studies it has been estimated that 35% of AD cases may be attributable to preventable causes such as low educational attainment, hearing loss, hypertension, obesity, smoking, depression, physical inactivity, social isolation and diabetes ^3^. However, lifestyle interventions that target modifiable risk factors are entirely dependent on accurate causal relationships being established between modifiable risk factors and AD. In observational studies, a correlation between a risk factor and AD cannot be reliably interpreted as evidence of a causal relationship due to potential confounding or reverse causation. Therefore, unless those modifiable factors specifically exacerbate disease progression, disease reduction strategies targeting them will not be successful.

Methods of causal inference that exploit genetic information, such as polygenic risk scores (PRS) and Mendelian randomization (MR), can overcome some of the limitations of observational studies. PRS are a measure of an individual’s genetic propensity to a trait and can be used in cross-trait analyses to test whether genetic liability for one trait is associated with disease risk for a second ^4^. While this does not imply that the trait causally modifies disease risk, since there are several alternative explanations, such a PRS-disease association would be expected if the trait were causal of disease, and thus PRS can be used to prioritize putative causal risk factors ^4^. MR uses genetic variants as proxies for environmental exposures to provide an estimate of the causal association between an intermediate exposure and a disease outcome. MR is akin to conducting a ‘genetic randomized control trial’, with the risk factors (genotypes) randomly allocated (from parents to offspring), independent of confounding factors that influence the risk factors and disease and unaffected by reverse causation ^5^. While MR can be used to directly assess causality between traits, it typically has lower statistical power than tests of PRS-disease associations ^4^.

In this study, we used PRS and MR to establish causal relationships between 22 modifiable risk factors and the AD phenome – AD status, AD age of onset survival (AAOS), CSF levels of amyloid-beta_42_ (Aβ_42_), tau and hyperphosphorylated tau (ptau_181_), hippocampal volume, cortical surface area and thickness, and the neuropathological burden of neuritic plaques, neurofibrillary tangles and vascular brain injury (VBI). Based on these analyses we identified a subset of modifiable risk factors that represent the most promising targets for public health initiatives to reduce AD burden in the population.

## Methods

### Genome-wide association summary statistics

We obtained GWAS summary statistics (GWAS-SS) for each exposure and outcome of interest (Table 1). Exposures included: alcohol consumption ^6^, the alcohol use disorder identification test (AUDIT) ^7^, moderate-vigorous physical activity (MVPA) ^8^, lipid traits ^9^, systolic blood pressure (SBP), diastolic blood pressure (DBP), pulse pressure (PP) ^10^, type 2 diabetes (T2D) ^11^, body mass index (BMI) ^12^, meat-related diet and a fish- and plant-related diet ^13^, depression ^14^, Insomnia symptoms ^15^, sleep duration ^16^, social isolation ^17^, smoking initiation ^6^, cigarettes per day ^6^, educational attainment ^18^, and hearing difficulty ^19^.

**Table 1:** GWAS datasets utilized in this study.

| Study | Trait | Cohort/Consortium | N | Age | Females (%) |
| --- | --- | --- | --- | --- | --- |
| **Exposures** |  |  |  |  |  |
| Liu et al 2019 | Alcohol Consumption | GSCAN; 23andMe | 941,280 | - | - |
|  | Smoking Initiation | GSCAN; 23andMe | 1,232,091 | - | - |
|  | Cigarettes per Day | GSCAN; 23andMe | 337,334 | - | - |
| Sanchez-Roige et al 2019 | Alcohol Use Disorder Test | UKBB; 23andMe | 141,932 | - | - |
| Wells et al 2019 | Hearing Difficulty | UKBB | 250,389 | - | - |
| Xue et al 2018 | Type 2 Diabetes | DIAGRAM; UKBB; GERA | 659,316 | - | - |
| Yengo et al 2018 | Body Mass Index | UKBB; GIANT | 690,495 | - | - |
| Willer et al 2013 | Total Cholesterol | GLC | 188,577 | 54.94 | 56.58 |
|  | LDL Cholesterol |  |  |  |  |
|  | HDL Cholesterol |  |  |  |  |
|  | Triglycerides |  |  |  |  |
| Evangelou et al 2018 | Diastolic Blood Pressure | UKBB; ICBP | 757,601 | - | - |
|  | Systolic Blood Pressure |  |  |  |  |
|  | Pulse Pressure |  |  |  |  |
| Howard et al 2019 | Depression | UKBB; PGC; deCODE; iPSYCH; GeneScotland; GERA; 23andMe | 807,553 | - | - |
| Jansen et al 2018 | Insomnia Symptoms | UKBB; 23andMe | 1,331,010 | - | - |
| Dashti et al 2019 | Sleep Duration | UKBB | 446,118 | 57.3 | 54.1 |
| Day et al 2018 | Social Isolation | UKBB | 452,302 | - | - |
| Lee et al 2018 | Educational Attainment | UKBB; SSGAC; 23andMe | 1,131,881 | 63.8 | 54.7 |
| Klimentidis et al 2018 | Moderate-Vigoures Physical Activity | UKBB | 377,234 | - | - |
| Niarchou et al 2020 | Meat-related diet | UKBB | 335,576 | - | 54% |
|  | Fish and plat related diet | UKBB | 335,576 | - | 54% |
| **Outcomes** |  |  |  |  |  |
| Lambert et al 2013 | Late Onset Alzheimer's disease | IGAP | 54,162 | 71 | 58.4 |
| Kunkle et al 2019 | Late Onset Alzheimer's disease | IGAP | 63,926 | 72.6 | 58.5 |
| Huang et al 2017 | Alzheimer's Age of Onset Survival | IGAP | 40,255 | 77.5 | 60.35 |
| Deming et al 2017 | CSF Aβ<sub>42</sub> | Knight-ADRC | 3,146 | 71.8 | 49.57 |
|  | CSF Ptau<sub>181</sub> |  |  |  |  |
|  | CSF Tau |  |  |  |  |
| Hibar et al 2015 | Hippocampal Volume | ENIGMA | 13,688 | 39.9 | 51.8 |
| Hibar et al 2017 | Hippocampal Volume | ENIGMA; CHARGE | 26,814 | 54.3 | 55.3 |
| Grasby et al 2020 | Cortical Surface Area | ENIGMA | 33,709 | 45.9 | 51.9 |
|  | Cortical Thickness |  |  |  |  |
| Beecham et al 2014 | Neuritic Plaques | ADGC | 4,914 | 74.7 | 65.4 |
|  | Neurofibrillary tangles |  |  |  |  |
|  | Vascular Brain Injury |  |  |  |  |

GWAS-SS for the AD phenome consisted of late-onset AD status ^20^, AAOS ^21^, CSF levels of Aβ_42_, ptau_181_ and total tau (Tau) ^22^, hippocampal volume ^23^, cortical surface area and thickness ^24^, neuropathological burden of neuritic plaques, neurofibrillary tangle burden and vascular brain injury ^25^. Due to data use restrictions associated with evaluating alcohol intake and education phenotypes in the most recent GWAS of AD and hippocampal volume, we used an earlier GWAS for AD ^26^ and hippocampal volume ^27^ for estimating the causal effect of alcohol intake and educational attainment on these phenotypes.

GWAS-SS that were mapped to earlier human genome builds were lifted over to Human Genome Build 19 ^28^. GWAS-SS were standardized using a pipeline, that 1) aligns effect alleles to the alternate allele on the forward strand of the human genome reference build and normalizes indels, 2) annotates variants with marker names using chromosome:position:ref:alt, 1000 Genomes rsIDs (phase 3), and dbSNP rsIDs (b151) 3) where allele frequencies are missing, annotates allele frequencies using non-Finnish Europeans from gnomAD (v2.1), and 4) convert summary statistics to VCF and TSV files.

### Alzheimer’s Disease Genetics Consortium

Individual-level genetic and phenotypic data used to compute and test the association of polygenic risk scores were obtained from the Alzheimer’s Disease Genetics Consortium (ADGC), a large multicenter project composed of 34 separate cohorts with the goal of performing genome-wide analyses of Alzheimer’s Disease. The recruitment and genotyping of ADGC samples has been described in detail elsewhere ^20,29^. Briefly, genotype data in each cohort underwent stringent quality control checks, with variants excluded if the call rate < 0.95, not in Hardy-Weinberg equilibrium (p < 1 × 10-6), and samples excluded if call rate was <0.95, discordant sex was reported based on X chromosome heterozygosity, cryptic relatedness, and non-European ancestry. Related individuals were determined within and across cohorts by identity-by-descent using KING ^30^, with individuals excluded based on a proportion of IBD < 0.1875, corresponding to less than halfway between second- and third-degree relatives. Ancestry was determined empirically by projecting samples onto principal components from known ancestral populations in the 1000 Genomes Project, with samples determined to be European population outliers if they were ±6 SD away from the EUR population mean on the first 10 principal components using PC-Air ^31^ and PLINK ^32^. SNPs that were not directly assayed were imputed on the Michigan Imputation Server individually for each of the cohorts or sub-cohorts using all ethnicities of the Haplotype Reference Consortium (HRC) 1.1 reference panel ^33^. Eagle was used for phasing and Minimac3 was used for imputation. Following imputation, poorly imputed (*r*^*2*^ < 0.8) or rare (MAF < 0.01) variants were removed and the cohorts merged for joint analysis. Following this merger, variants with low call rate due to differential imputation (< 95%) were removed, and then samples with low call rate (< 95%) were removed. Within-ancestry principal components were created using PLINK to correct for residual population stratification within the European population subset. After sample QC, 26,431 participants were available (Table 2). Written informed consent was obtained from study participants or, for those with substantial cognitive impairment, from a caregiver, legal guardian, or other proxy, and the study protocols for all populations were reviewed and approved by the appropriate Institutional review boards (IRB’s).

**Table 2:**
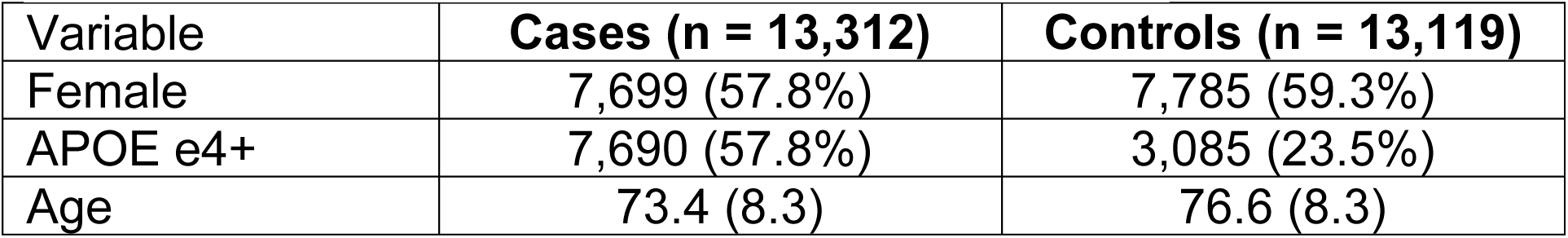
Demographic characteristics of ADGC.

| Variable | Cases (n = 13,312) | Controls (n = 13,119) |
| --- | --- | --- |
| Female | 7,699 (57.8%) | 7,785 (59.3%) |
| APOE e4+ | 7,690 (57.8%) | 3,085 (23.5%) |
| Age | 73.4 (8.3) | 76.6 (8.3) |

### Polygenic Risk Scores

The software package *PRSice-2* was used to construct polygenic risk scores for each of the exposures of interest in ADGC ^34^. *PRSice* generates PRS as the sum of all alleles associated with the exposure of interest exceeding a given *P*-value threshold (*P*_*t*_), weighted by their effect size estimated in an independent GWAS on the trait. SNPs were clumped to obtain variants in linkage equilibrium with an *r*^*2*^ > 0.001 within a 10MB window and PRS were constructed across a range of *P*_*t*_ (*P*_*t*_ = 5e-8, 1e-6, 1e-5, 1e-4, 1e-3, 0.01, 0.05, 0.1, 0.2, 0.3, 0.4, and 0.5). The optimal *P*-value threshold was determined according to the results of a linear regression (PRSice uses linear regression with binary traits to avoid issues of perfect separation during permutation) testing the association of the trait PRS and the AD outcome, adjusted for age, sex, *APOE* ε4 dose, and 10 principal components; the PRS *P*_*t*_ with the smallest *P*-value of association is selected for association analysis. To guard against overfitting, 1000 permutations were conducted to obtain an empirical *P*-value for each PRS-AD association. PRS were standardized to have a mean of 0 and SD of 1. After obtaining the optimal *P*_t_ the association between each exposure PRS and AD was evaluated using logistic regression adjusting for age, sex, *APOE* ε4 dose, and 10 principal components. The Benjamini & Hochberg false discovery rate was used to account for the multiple testing across the 22 different exposures.

### Mendelian Randomization Analysis

#### Genetic Instruments

For each exposure, we constructed two different sets of instrumental variables (IV), corresponding to independent (1) genome-wide significant SNPs *(P* < 5 × 10^−8^) and (2) SNPs of at least borderline significance *(P* < 5 × 10^−6^). Increasing the number of SNPs used as IVs increases the phenotypic variance explained and, thus, has the potential to increase statistical power. However, if the additional variants included violate the core MR assumptions then they may instead reduce power, biasing the results towards the null by introducing weak instrument bias. To obtain independent SNPs, linkage disequilibrium (LD) clumping was performed by excluding SNPs that have an r2 > 0.001 with another variant with a smaller p-value association within a 10MB window using PLINK ^32^. For genetic variants that were not present in the outcome GWAS, PLINK was used to identify proxy SNPs that were in LD (r^2^ > 0.8; EUR reference population). Finally, the exposure and outcome GWAS datasets were harmonized so that the effect size for the exposure and outcome corresponded to the same effect alleles. Genetic variants that were palindromic with ambiguous allele frequencies (AF > 0.42), or that had incompatible alleles, were removed. Variants within the *APOE* region were excluded due to pleiotropy with AD. The proportion of variance in the phenotype explained by each instrument and F-statistic were calculated as previously described ^35,36^.

### Statistical Analysis

For each genetic variant, we calculated an instrumental variable ratio estimate by dividing the SNP-exposure by SNP-outcome and the resulting coefficients were combined in a fixed-effects meta-analysis using an inverse-variance weighted (IVW) approach to give an overall estimate of causal effect ^5^. The IVW method assumes that all SNPs included in the causal estimate are valid instruments - that is, that they do not violate any of the underlying MR assumptions, in particular horizontal pleiotropy, whereby genetic variants have direct effects on multiple phenotypes, could lead to false inference of causal associations ^5^. In order to account for potential violations of the assumptions underlying the IVW analysis, we conducted sensitivity analyses using alternative MR methods known to be more robust to horizontal pleiotropy in particular, but at the cost of reduced statistical power. The alternative approaches included 1) Weighted Median Estimator (WME), which tests the median effect of all of the IV variants, allowing 50% of variants to exhibit horizontal pleiotropy ^5^; 2) Weighted Mode Based Estimator (WMBE), which clusters variants into groups based on the similarity of causal effects and reports the final causal effect based on the cluster with the largest number of variants ^5^; and 3) MR-Egger regression, which allows all variants to be subject to direct effects that bias the estimate in the same direction ^5^.

The MR-Egger regression intercept was used to verify the absence of pleiotropic effects of the SNPs on the outcome ^5^. To further confirm the absence of distortions in the causal effects due to heterogeneity or horizontal pleiotropy, we used the Mendelian randomization pleiotropy residual sum and outlier (MR-PRESSO) test to detect and correct for horizontal pleiotropic outliers ^37^. Where heterogeneity was detected (the MR-PRESSO Global Test) and significant outliers were detected (MR-PRESSO Outlier Test), the outliers were removed.

We report the IVW results for the set of IV variants (*at P* < 1×10^−8^ or 1×10^−6^) with the smallest p-value, outliers were removed if detected. Where there was evidence of horizontal pleiotropy or heterogeneity (MR-PRESSO Global Test *p* < 0.05 or an MR-Egger Intercept *p* < 0.05), we report the IVW results for which the sensitivity analyses were also significant and the effect direction was concordant with the IVW results. To account for multiple testing, we report q-values, a false discovery rate-based measure of significance ^38^. Power analyses were conducted using the non-centrality parameter-based approach using the observed IVW coefficient ^39^.

All statistical analyses were conducted using R version 3.5.2. Mendelian randomization analysis was performed using the ‘TwoSampleMR’ package ^5^. A Snakemake workflow was constructed that automates the PRS and MR analysis pipelines and allows for multiple exposure – outcomes datasets to be run in parallel ^40^.

The SNPs used as IVs, their harmonized effects and outliers are presented in Supplementary Table 1. The causal estimates for each p-value threshold, MR method and pre- and post-outlier removal are presented in Supplementary Table 2.

## Results

### Polygenic Risk Score Analysis

We evaluated the association of 22 PRS for potentially modifiable risk factors with AD in ADGC. The P_t_, number of SNPs and the association of each PRS with AD are presented in Table 3. After correction for multiple testing, a 1SD higher PRS for educational attainment increased risk of AD (OR [CI] 0.93 [0.91, 0.96]). Higher PRS for total cholesterol levels (OR [CI]1.05 [1.02, 1.08]) and moderate-vigorous physical activity (OR [CI]1.04 [1.01, 1.07]) were associated with an increased risk of AD. Using only genome-wide significant SNPs, only the PRS for educational attainment was significant after correction for multiple testing (Table 3).

**Table 3:**
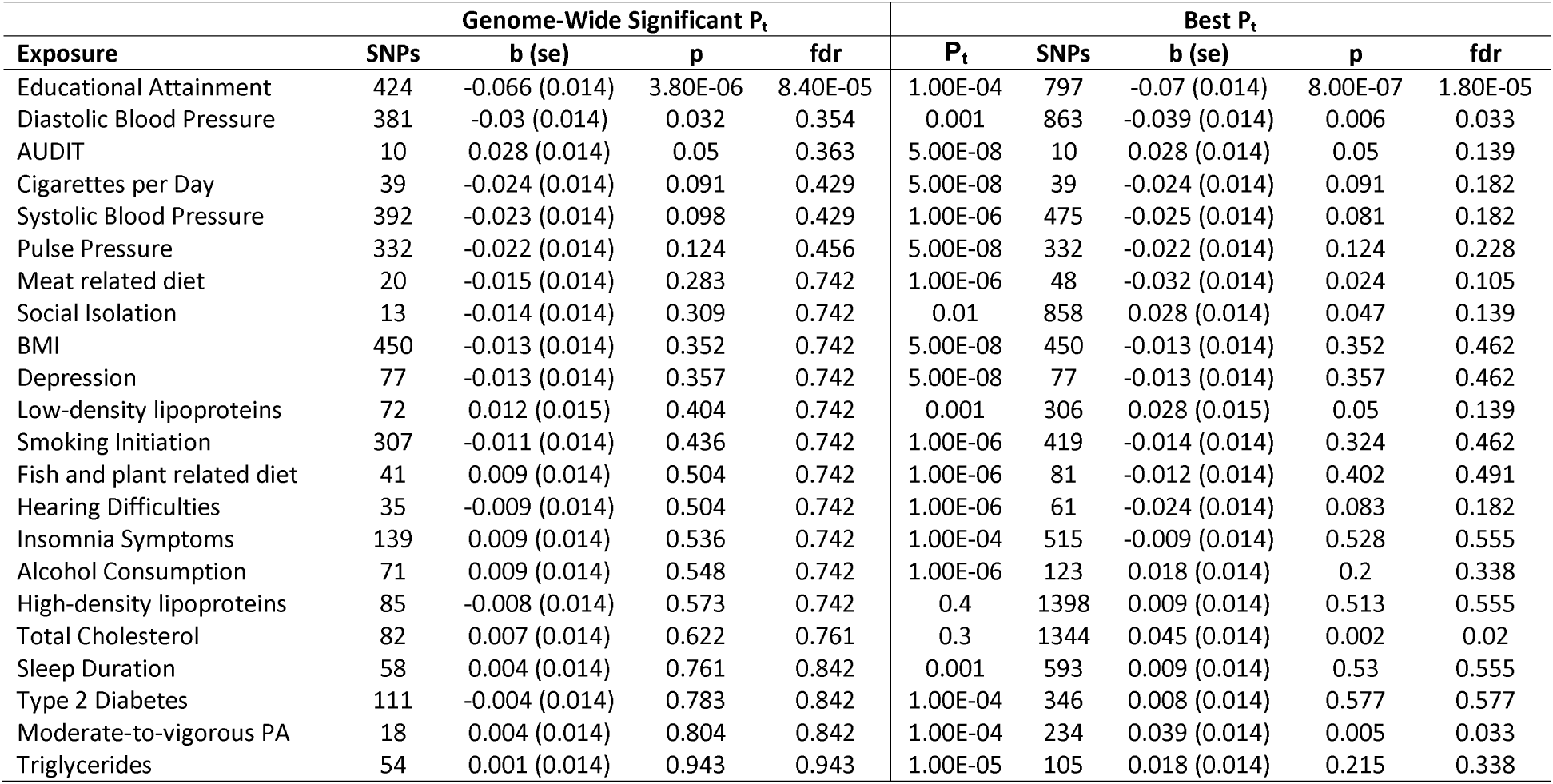
Association of polygenic risk scores for potentially modifiable risk factors on Alzheimer’s disease.

| Exposure | Genome-Wide Significant $P_t$ | | | | Best $P_t$ | | | | |
| --- | --- | --- | --- | --- | --- | --- | --- | --- | --- |
| | SNPs | b (se) | p | fdr | $P_t$ | SNPs | b (se) | p | fdr |
| Educational Attainment | 424 | -0.066 (0.014) | 3.80E-06 | 8.40E-05 | 1.00E-04 | 797 | -0.07 (0.014) | 8.00E-07 | 1.80E-05 |
| Diastolic Blood Pressure | 381 | -0.03 (0.014) | 0.032 | 0.354 | 0.001 | 863 | -0.039 (0.014) | 0.006 | 0.033 |
| AUDIT | 10 | 0.028 (0.014) | 0.05 | 0.363 | 5.00E-08 | 10 | 0.028 (0.014) | 0.05 | 0.139 |
| Cigarettes per Day | 39 | -0.024 (0.014) | 0.091 | 0.429 | 5.00E-08 | 39 | -0.024 (0.014) | 0.091 | 0.182 |
| Systolic Blood Pressure | 392 | -0.023 (0.014) | 0.098 | 0.429 | 1.00E-06 | 475 | -0.025 (0.014) | 0.081 | 0.182 |
| Pulse Pressure | 332 | -0.022 (0.014) | 0.124 | 0.456 | 5.00E-08 | 332 | -0.022 (0.014) | 0.124 | 0.228 |
| Meat related diet | 20 | -0.015 (0.014) | 0.283 | 0.742 | 1.00E-06 | 48 | -0.032 (0.014) | 0.024 | 0.105 |
| Social Isolation | 13 | -0.014 (0.014) | 0.309 | 0.742 | 0.01 | 858 | 0.028 (0.014) | 0.047 | 0.139 |
| BMI | 450 | -0.013 (0.014) | 0.352 | 0.742 | 5.00E-08 | 450 | -0.013 (0.014) | 0.352 | 0.462 |
| Depression | 77 | -0.013 (0.014) | 0.357 | 0.742 | 5.00E-08 | 77 | -0.013 (0.014) | 0.357 | 0.462 |
| Low-density lipoproteins | 72 | 0.012 (0.015) | 0.404 | 0.742 | 0.001 | 306 | 0.028 (0.015) | 0.05 | 0.139 |
| Smoking Initiation | 307 | -0.011 (0.014) | 0.436 | 0.742 | 1.00E-06 | 419 | -0.014 (0.014) | 0.324 | 0.462 |
| Fish and plant related diet | 41 | 0.009 (0.014) | 0.504 | 0.742 | 1.00E-06 | 81 | -0.012 (0.014) | 0.402 | 0.491 |
| Hearing Difficulties | 35 | -0.009 (0.014) | 0.504 | 0.742 | 1.00E-06 | 61 | -0.024 (0.014) | 0.083 | 0.182 |
| Insomnia Symptoms | 139 | 0.009 (0.014) | 0.536 | 0.742 | 1.00E-04 | 515 | -0.009 (0.014) | 0.528 | 0.555 |
| Alcohol Consumption | 71 | 0.009 (0.014) | 0.548 | 0.742 | 1.00E-06 | 123 | 0.018 (0.014) | 0.2 | 0.338 |
| High-density lipoproteins | 85 | -0.008 (0.014) | 0.573 | 0.742 | 0.4 | 1398 | 0.009 (0.014) | 0.513 | 0.555 |
| Total Cholesterol | 82 | 0.007 (0.014) | 0.622 | 0.761 | 0.3 | 1344 | 0.045 (0.014) | 0.002 | 0.02 |
| Sleep Duration | 58 | 0.004 (0.014) | 0.761 | 0.842 | 0.001 | 593 | 0.009 (0.014) | 0.53 | 0.555 |
| Type 2 Diabetes | 111 | -0.004 (0.014) | 0.783 | 0.842 | 1.00E-04 | 346 | 0.008 (0.014) | 0.577 | 0.577 |
| Moderate-to-vigorous PA | 18 | 0.004 (0.014) | 0.804 | 0.842 | 1.00E-04 | 234 | 0.039 (0.014) | 0.005 | 0.033 |
| Triglycerides | 54 | 0.001 (0.014) | 0.943 | 0.943 | 1.00E-05 | 105 | 0.018 (0.014) | 0.215 | 0.338 |

### Mendelian Randomization Analysis

We used Mendelian randomization to estimate the causal associations between 22 potentially modifiable risk factors and 11 AD outcomes, across two sets of IV variants corresponding to two different p-value thresholds. We observed 12 exposure-outcome pairs that were significant at an FDR < 0.05 and that either showed no evidence of heterogeneity or horizontal pleiotropy, or in the presence of heterogeneity or horizontal pleiotropy, the additional MR sensitivity analyses were significant (Figure 1; Table 4). The PVE, F-statistics and power for each model are presented in Supplementary Table 2.

**Table 4:**
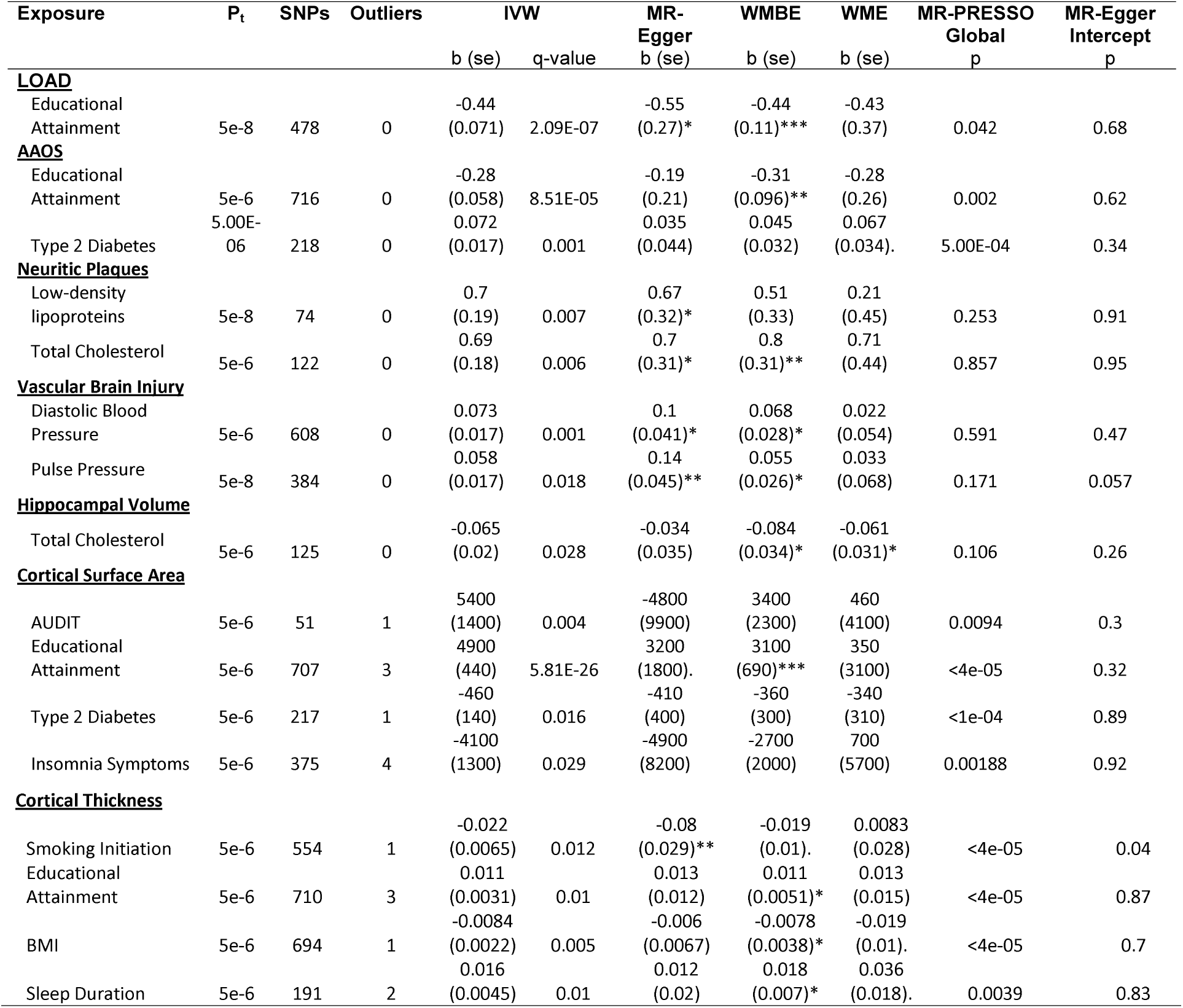
Causal association of potentially modifiable risk factors on Alzheimer’s disease and Alzheimer’s endophenotypes.

**Figure 1:**
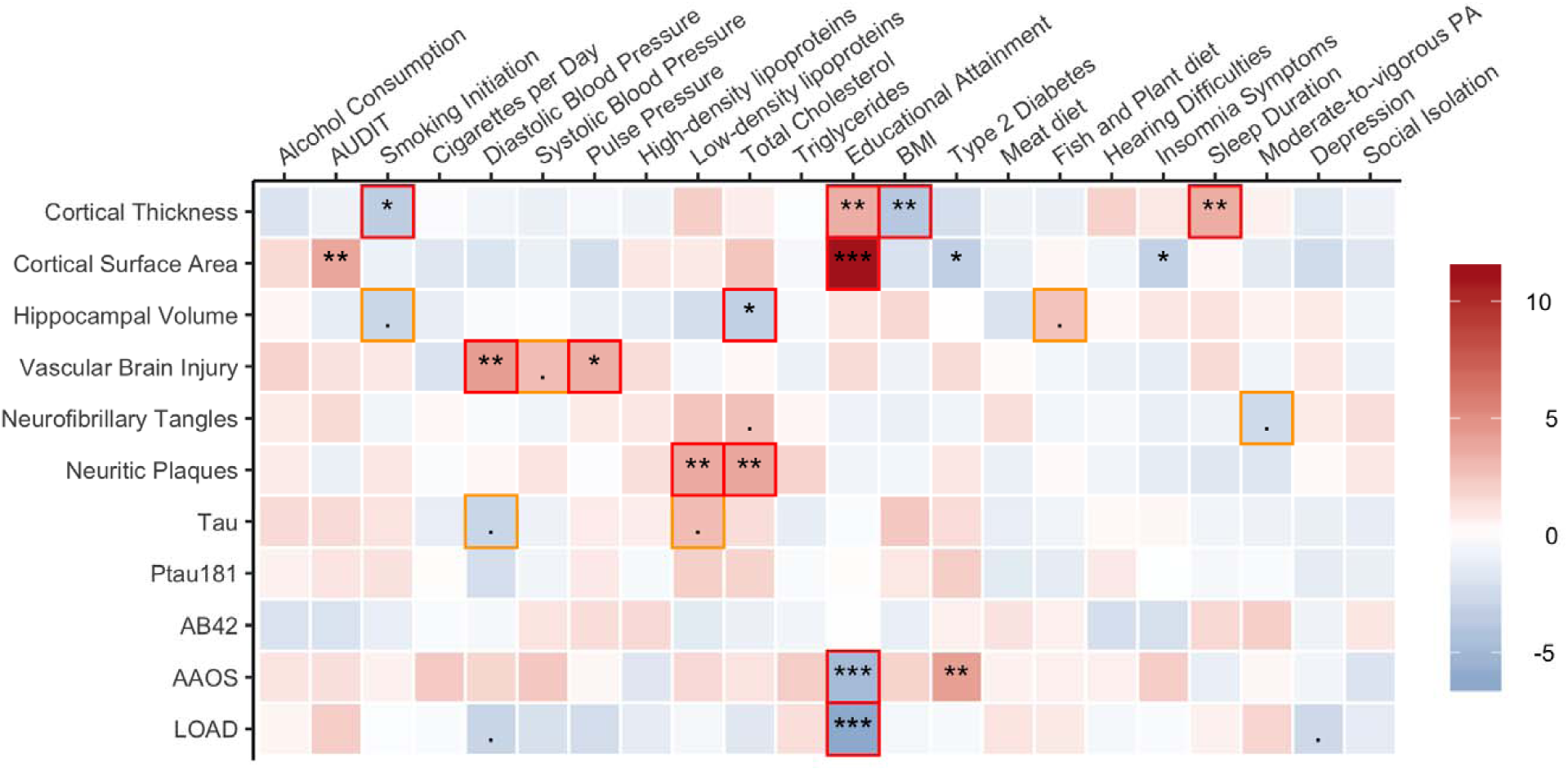
Putative causal associations between modifiable risk factors and the AD phenome. Shown are the best IVW results for each causal association, with colors representing the standardized effect sizes - for LOAD, NP, NFT, and AAOS red indicates increased risk / earlier onset and blue reduced risk / delayed onset, for CSF levels and Hippocampal volume, red indicates increased levels/volume and blue reduced levels/volume. “.” FDR < 0.1; * FDR < 0.05; ** FDR < 0.01; *** FDR < 0.001. Causal estimates bracketed in red or orange indicate significant causal effects that showed no evidence for horizontal pleiotropy or where sensitivity analyses were also significant.

Genetically predicted increased low-density lipoproteins (OR [CI]: 2.01 [1.39, 2.92]) and total cholesterol levels (OR [CI]: 1.99 [1.4, 2.84]) were associated with significantly increased risk of neuritic plaques. Genetically predicted higher diastolic blood pressure (OR [CI]: 1.08 [1.04, 1.11]) and pulse pressure (OR [CI]: 1.06 [1.02, 1.1]) were associated with significantly increased risk of vascular brain injury. Genetically predicted higher educational attainment was associated with significantly 1) lower risk of Alzheimer’s disease (OR [CI]: 0.64 [0.56, 0.74]), 2) delayed AAOS (HR [CI]: 0.76 [0.67, 0.85]), 3) increased cortical surface area (β mm^2^ [CI]: 4900 [4037.6, 5762.4]), and 4) increased cortical thickness (β mm [CI]: 0.01 [0, 0.02]). Genetically predicted longer sleep duration was associated with significantly increased cortical thickness after outlier removal (β mm [CI]: 0.02 [0.01, 0.02]). Genetically predicted smoking status was associated with significantly reduced cortical thickness (β mm [CI]: -0.02 [-0.03, -0.01]). Genetically predicted higher BMI was associated with significantly reduced cortical thickness after outlier removal (β mm [CI]: -0.01 [-0.01, 0]).

A further three risk factors, including AUDIT, diabetes, and insomnia, were causally associated with the AD phenome in the IVW analysis (Table 4; Figure 1), however, there was evidence of heterogeneity and the sensitivity analyses were non-significant suggesting that the observed associations were not robust to violations of MR underlying assumptions.

## Discussion

Using genetic variants as proxies for modifiable risk factors, we applied PRS and MR analyses to investigate the association of putative modifiable risk factors with the AD phenome. PRS for higher educational attainment and diastolic blood pressure were observed to be associated with reduced risk for AD, while higher total cholesterol and increased moderate-vigorous physical activity were associated with an increased risk of AD. However, in the MR analysis, only higher educational attainment was causally associated with a reduced risk of AD. The lack of causal associations between modifiable risk factors and AD may reflect heterogeneity in the underlying pathogenesis that can lead to clinical phenotypes analogous to Alzheimer’s disease.

An endophenotype is usually less genetically complex than the disorder it underlies due to the endophenotype being influenced by fewer genetic risk factors than the disease as a whole and reflecting a single pathophysiological pathway of the overall clinical disorder. As endophenotypes can be measured in both cases and controls there is greater power to detect an association due to the effect allele influencing the endophenotype even in asymptomatic carriers. As such we expanded our MR analysis to evaluate the causal effect of modifiable risk factors on AD endophenotypes to evaluate how potential risk factors may influence the underlying pathophysiological pathways of AD. We observed 1) higher total-cholesterol and LDL-cholesterol levels to be causally associated with increased risk of neuritic plaque burden, 2) higher diastolic blood pressure and pulse pressure causally associated with increased risk of vascular brain injury, and 3) higher educational attainment causally associated with a delayed AAOS and increased cortical surface area and thickness. Furthermore, 1) higher total cholesterol was causally associated with decreased hippocampal volume, 2) smoking status and higher BMI were causally associated with reduced cortical thickness, and 3) longer sleep duration was causally associated with increased cortical thickness.

Observational studies have indicated that lifestyle interventions targeting modifiable risk factors can either prevent or delay the age of onset of dementia. In particular, low educational attainment, hearing loss, hypertension, obesity, smoking, depression, physical inactivity, social isolation and diabetes have been indicated to be key risk factors in the development of dementia ^3^. However, with the exception of educational attainment, our analyses did not provide strong evidence of a causal association with these risk factors and AD or AAOS. The lack of a causal association between these risk factors and AD could be due to insufficient power in our analyses, but, alternatively, may be a result of confounding or reverse causation in observational studies. For instance, increased physical activity is generally associated with a reduced risk of dementia ^3^, however, a recent meta-analysis found that the protective association with dementia was observed when physical activity was measured <10 years before dementia diagnosis, but when measured >10 years before dementia onset no association with dementia was observed – consistent with reverse causation driving the observed protective association ^41^. Additionally, while these risk factors may not be associated with AD pathogenesis, they may be associated with the pathogenesis of other dementia subtypes. For instance, the observed association between blood pressure and VBI suggests that while reducing blood pressure in late life may have limited utility in the prevention of AD, it may reduce the risk of vascular dementia by reducing the risk of VBI and therefore affect the risk for all-cause dementia, but not specifically affect the risk of AD.

The association of modifiable risk factor PRS with clinically diagnosed AD has not been extensively studied, though several studies have conducted phenome-wide scans to evaluate the association of AD PRS with a wide range of diseases and other traits. Using data from the UK Biobank (n = 334,398), Richardson and colleagues found that an AD PRS composed of 124 SNPs and inclusive of *APOE* (Pt ≤ 5e-05) was associated with 72/551 traits (FDR < 0.05) ^4^. In particular, a higher AD PRS was associated with lower diastolic blood pressure and BMI, reduced risk of self-reported diabetes, shorter sleep duration, increased risk of self-reported high cholesterol and increased amount of moderate-physical activity ^4^. Similarly, a second study by Korologou-Linden and colleagues evaluated the association of an AD PRS composed of 18 SNPs, inclusive of *APOE*, (Pt ≤ 5e-08) across 15,403 traits in the UK Biobank (n = 334,968) ^42^. A higher AD PRS was associated with 165 traits and in particular, with lower diastolic blood pressure, lower BMI, increased total cholesterol, levels, reduced risk of self-reported diabetes, increased oily fish consumption, increased sleeplessness or insomnia, reduced sleep duration increased amount of moderate-physical activity and increased risk of self-reported depression ^42^. In a follow-up MR analysis of these traits, only moderate-physical activity was observed to be causally associated with an increased risk of AD ^42^.

Two earlier studies used MR to evaluate the association of potentially modifiable risk with AD cases-control status ^43,44^. First Østergaard and colleagues evaluated the association of 13 risk factors with AD and observed that higher systolic blood pressure, HDL-cholesterol and smoking quantity were associated with a reduced risk of AD, while higher total cholesterol and LDL cholesterol were associated with increased risk ^44^. No significant associations were observed for BMI, diabetes, insulin resistance, triglycerides, smoking initiation or education and after variants in the *APOE* locus were excluded from the analysis, the cholesterol levels were no longer significantly associated with AD risk ^44^. Second, Larsson and colleagues evaluated the association of 22 risk factors with AD, finding that years of education, intelligence, and 25-hydroxyvitamin D were associated with a reduced risk of AD, while coffee consumption was associated with increased risk ^43^. No significant associations were observed between alcohol consumption, serum folate, serum vitamin B_12_, homocysteine, cardiometabolic factors or C reactive protein with Alzheimer’s disease ^43^.

The results of this study should be interpreted in conjunction with knowledge of its limitations and those of MR in general. First, while we cannot exclude that our findings may be affected by weak instrument bias, the F-statistics for all of the analyses were greater than 10, indicating that the instrument strength was sufficient for MR analysis ^36^. However, in two-sample MR, weak instrument bias is in the direction of the null, thus, we cannot exclude low power as an explanation for the null results ^45^. Second, we cannot completely rule out violations of the independence and the exclusion restriction assumption, particularly in regard to pleiotropy ^46^. Nevertheless, we used several methods to identify robust causal estimates, including outlier removal using MR-PRESSO and WMBE, WME and MR-Egger sensitivity analyses. Finally, it is assumed that both samples used to generate the GWAS summary statistics used in the MR model come from comparable populations. In evaluating the demographics of the studies used in this analysis, the exposures have an average age of 56.1 – 63.8yrs, while outcomes, with the exception of hippocampal volume, have an average age of 71 – 74.7yrs. As such, some of the results reported here may be subject to survivor bias ^47^. Nevertheless, the bias introduced by survival effects is large for exposures that strongly affect survival. However, when selection effects are weak or moderate, selection bias does not adversely affect causal estimates ^47^.

Despite these limitations, this study has significant strengths. We assessed the causal effect of multiple modifiable factors strongly hypothesized as affecting AD risk. In addition, we used the largest GWAS for AD and the exposure traits available at the time of analysis, allowing us to include the largest possible number of instruments for the exposures, resulting in increased statistical power. Finally, rather than limiting our analyses to AD case/control status, we expanded our MR analysis to include AD endophenotypes.

In conclusion, this study used large exposure and outcome GWAS to conduct PRS and MR analyses to evaluate the causal association of potentially modifiable risk factors with the AD phenome. The PRS analysis identified four traits for which a higher genetic predisposition influenced AD risk. In the follow-up MR analysis, only genetically predicted higher education was observed to have a causal association with reduced AD risk. Expanding our analysis to additional AD endophenotypes, we observed that higher genetically predicted cholesterol levels and blood pressure were associated with increased risk of neuritic plaque burden and vascular brain injury respectively, suggesting that these risk factors influence the development of neurodegenerative disease pathology.

## Acknowledgments

SJA, BFH, EM and AMG were supported by the JPB Foundation (http://www.jpbfoundation.org) and by the National Institute of Health (U01AG052411 and U01AG058635; principal investigator Alison Goate). PFO was supported by funding from the UK Medical Research Council (MR/N015746/1) and the National Institute of Health (R01MH122866). Adam Naj was supported by funding from the National Institute of Aging (R01 AG054060 and RF1 AG061351). The funders had no role in study design, data collection and analysis, decision to publish, or preparation of the manuscript. This analysis was possible due to the generous sharing of genome-wide association summary statistics. We would like to thank the research participants and employees of 23andMe for making this work possible.

ADGC: The Alzheimer’s Disease Genetics Consortium supported collection and genotyping of samples used in this study through National Institute on Aging (NIA) grants U01AG032984 and RC2AG036528.

NCRAD: Samples from the National Centralized Repository for Alzheimer’s Disease and Related Dementias (NCRAD), which receives government support under a cooperative agreement grant (U24 AG21886) awarded by the National Institute on Aging (NIA), were used in this study. We thank contributors who collected samples used in this study, as well as patients and their families, whose help and participation made this work possible.

NACC: The NACC database is funded by NIA/NIH Grant U01 AG016976. NACC data are contributed by the NIA-funded ADCs: P30 AG019610 (PI Eric Reiman, MD), P30 AG013846 (PI Neil Kowall, MD), P30 AG062428-01 (PI James Leverenz, MD) P50 AG008702 (PI Scott Small, MD), P50 AG025688 (PI Allan Levey, MD, PhD), P50 AG047266 (PI Todd Golde, MD, PhD), P30 AG010133 (PI Andrew Saykin, PsyD), P50 AG005146 (PI Marilyn Albert, PhD), P30 AG062421-01 (PI Bradley Hyman, MD, PhD), P30 AG062422-01 (PI Ronald Petersen, MD, PhD), P50 AG005138 (PI Mary Sano, PhD), P30 AG008051 (PI Thomas Wisniewski, MD), P30 AG013854 (PI Robert Vassar, PhD), P30 AG008017 (PI Jeffrey Kaye, MD), P30 AG010161 (PI David Bennett, MD), P50 AG047366 (PI Victor Henderson, MD, MS), P30 AG010129 (PI Charles DeCarli, MD), P50 AG016573 (PI Frank LaFerla, PhD), P30 AG062429-01(PI James Brewer, MD, PhD), P50 AG023501 (PI Bruce Miller, MD), P30 AG035982 (PI Russell Swerdlow, MD), P30 AG028383 (PI Linda Van Eldik, PhD), P30 AG053760 (PI Henry Paulson, MD, PhD), P30 AG010124 (PI John Trojanowski, MD, PhD), P50 AG005133 (PI Oscar Lopez, MD), P50 AG005142 (PI Helena Chui, MD), P30 AG012300 (PI Roger Rosenberg, MD), P30 AG049638 (PI Suzanne Craft, PhD), P50 AG005136 (PI Thomas Grabowski, MD), P30 AG062715-01 (PI Sanjay Asthana, MD, FRCP), P50 AG005681 (PI John Morris, MD), P50 AG047270 (PI Stephen Strittmatter, MD, PhD).

NIAGADS: Data for this study were prepared, archived, and distributed by the National Institute on Aging Alzheimer’s Disease Data Storage Site (NIAGADS) at the University of Pennsylvania (U24 AG041689).

## Author Contributions

SJA, AMG, PFO, BFH, EM contributed to the conception and design of the study. SJA, BFH, EM. contributed to the acquisition and analysis of data. SJA, AMG, PFO, BFH, EM. contributed to drafting a significant portion of the manuscript or figures. LAF, JLH, RM, ACN, MAPV, GDS, LW contributed to the acquisition of data for the Alzheimer’s Disease Genetics Consortium.

## Data Availability

This study used published summary results from published research papers, with the references for those studies provided in the main paper. Supplementary Table 1 provides the harmonized SNP effects needed to reproduce the results of this analysis.

## Supplementary Data

Supplementary Table 1: Harmonized SNP effects across exposures – outcomes

Supplementary Table 2: Mendelian Randomization results

## References

1. 2020 Alzheimer’s disease facts and figures. Alzheimer’s Dementia 2020;16(3):391–460.

2. Anstey KJ, Ee N, Eramudugolla R, et al. A Systematic Review of Meta-Analyses that Evaluate Risk Factors for Dementia to Evaluate the Quantity, Quality, and Global Representativeness of Evidence. J Alzheimer’s Dis Jad 2019;1–21.

3. Livingston G, Sommerlad A, Orgeta V, et al. Dementia prevention, intervention, and care. Lancet 2017;390(10113):2673–2734.

4. Richardson TG, Harrison S, Hemani G, Smith GD. An atlas of polygenic risk score associations to highlight putative causal relationships across the human phenome. Elife 2019;8:e43657.

5. Hemani G, Zheng J, Elsworth B, et al. The MR-Base platform supports systematic causal inference across the human phenome. Elife 2018;7:e34408.

6. Liu M, Jiang Y, Wedow R, et al. Association studies of up to 1.2 million individuals yield new insights into the genetic etiology of tobacco and alcohol use. Nat Genet 2019;51(2):237–244.

7. Sanchez-Roige S, Palmer AA, Fontanillas P, et al. Genome-Wide Association Study Meta-Analysis of the Alcohol Use Disorders Identification Test (AUDIT) in Two Population-Based Cohorts. Am J Psychiat 2019;176(2):107–118.

8. Klimentidis YC, Raichlen DA, Bea J, et al. Genome-wide association study of habitual physical activity in over 377,000 UK Biobank participants identifies multiple variants including CADM2 and APOE. Int J Obes 2005 2018;42(6):1161–1176.

9. Willer CJ, Schmidt EM, Sengupta S, et al. Discovery and refinement of loci associated with lipid levels. Nat Genet 2013;45(11):1274–83.

10. Evangelou E, Warren HR, Mosen-Ansorena D, et al. Genetic analysis of over 1 million people identifies 535 new loci associated with blood pressure traits. Nat Genet 2018;50(10):1412–1425.

11. Xue A, Wu Y, Zhu Z, et al. Genome-wide association analyses identify 143 risk variants and putative regulatory mechanisms for type 2 diabetes. Nat Commun 2018;9(1):2941.

12. Yengo L, Sidorenko J, Kemper KE, et al. Meta-analysis of genome-wide association studies for height and body mass index in □700000 individuals of European ancestry. Hum Mol Genet 2018;27(20):3641–3649.

13. Niarchou M, Byrne EM, Trzaskowski M, et al. Genome-wide association study of dietary intake in the UK biobank study and its associations with schizophrenia and other traits. Transl Psychiat 2020;10(1):51.

14. Howard DM, Adams MJ, Clarke T-K, et al. Genome-wide meta-analysis of depression identifies 102 independent variants and highlights the importance of the prefrontal brain regions. Nat Neurosci 2019;22(3):343–352.

15. Jansen PR, Watanabe K, Stringer S, et al. Genome-wide analysis of insomnia in 1,331,010 individuals identifies new risk loci and functional pathways. Nat Genet 2019;51(3):394–403.

16. Dashti HS, Jones SE, Wood AR, et al. Genome-wide association study identifies genetic loci for self-reported habitual sleep duration supported by accelerometer-derived estimates. 2019;

17. Day FR, Ong KK, Perry JRB. Elucidating the genetic basis of social interaction and isolation. Nat Commun 2018;9(1):2457.

18. Lee JJ, Wedow R, Okbay A, et al. Gene discovery and polygenic prediction from a genome-wide association study of educational attainment in 1.1 million individuals. Nat Genet 2018;50(8):1112–1121.

19. Wells HRR, Freidin MB, Abidin FNZ, et al. GWAS Identifies 44 Independent Associated Genomic Loci for Self-Reported Adult Hearing Difficulty in UK Biobank. Am J Hum Genetics 2019;105(4):788–802.

20. Kunkle BW, Grenier-Boley B, Sims R, et al. Genetic meta-analysis of diagnosed Alzheimer’s disease identifies new risk loci and implicates Aβ, tau, immunity and lipid processing. Nat Genet 2019;51(3):414–430.

21. Huang K-L, Marcora E, Pimenova AA, et al. A common haplotype lowers PU.1 expression in myeloid cells and delays onset of Alzheimer’s disease. Nat Neurosci 2017;20(8):1052–1061.

22. Deming Y, Li Z, Kapoor M, et al. Genome-wide association study identifies four novel loci associated with Alzheimer’s endophenotypes and disease modifiers. Acta Neuropathol 2017;133(5):839–856.

23. Hibar DP, Adams HHH, Jahanshad N, et al. Novel genetic loci associated with hippocampal volume. Nat Commun 2017;8(1):13624.

24. Grasby KL, Jahanshad N, Painter JN, et al. The genetic architecture of the human cerebral cortex. Science 2020;367(6484):eaay6690.

25. Beecham GW, Hamilton K, Naj AC, et al. Genome-Wide Association Meta-analysis of Neuropathologic Features of Alzheimer’s Disease and Related Dementias. Plos Genet 2014;10(9):e1004606.

26. Lambert J-C, Ibrahim-Verbaas CA, Harold D, et al. Meta-analysis of 74,046 individuals identifies 11 new susceptibility loci for Alzheimer’s disease. Nat Genet 2013;45(12):1452–1458.

27. Hibar DP, Stein JL, Renteria ME, et al. Common genetic variants influence human subcortical brain structures. Nature 2015;520(7546):224–229.

28. Kent WJ, Sugnet CW, Furey TS, et al. The Human Genome Browser at UCSC. Genome Res 2002;12(6):996–1006.

29. Naj AC, Jun G, Beecham GW, et al. Common variants at MS4A4/MS4A6E, CD2AP, CD33 and EPHA1 are associated with late-onset Alzheimer’s disease. Nat Genet 2011;43(5):436–441.

30. Manichaikul A, Mychaleckyj JC, Rich SS, et al. Robust relationship inference in genome-wide association studies. Bioinformatics 2010;26(22):2867–2873.

31. Conomos MP, Miller MB, Thornton TA. Robust inference of population structure for ancestry prediction and correction of stratification in the presence of relatedness. Genet Epidemiol 2015;39(4):276–93.

32. Purcell S, Neale B, Todd-Brown K, et al. PLINK: A Tool Set for Whole-Genome Association and Population-Based Linkage Analyses. Am J Hum Genetics 2007;81(3):559–575.

33. Consortium HR, McCarthy S, Das S, et al. A reference panel of 64,976 haplotypes for genotype imputation. Nat Genet 2016;48(10):ng.3643.

34. Choi SW, O’Reilly PF. PRSice-2: Polygenic Risk Score software for biobank-scale data. Gigascience 2019;8(7)

35. Shim H, Chasman DI, Smith JD, et al. A multivariate genome-wide association analysis of 10 LDL subfractions, and their response to statin treatment, in 1868 Caucasians. Plos One 2015;10(4):e0120758.

36. Burgess S, Thompson SG, Collaboration CCG. Avoiding bias from weak instruments in Mendelian randomization studies. Int J Epidemiol 2011;40(3):755–764.

37. Verbanck M, Chen C-Y, Neale B, Do R. Detection of widespread horizontal pleiotropy in causal relationships inferred from Mendelian randomization between complex traits and diseases. Nat Genet 2018;50(5):693–698.

38. Storey JD. A direct approach to false discovery rates. J Royal Statistical Soc Ser B Statistical Methodol 2002;64(3):479–498.

39. Brion M-JA, Shakhbazov K, Visscher PM. Calculating statistical power in Mendelian randomization studies. Int J Epidemiol 2013;42(5):1497–501.

40. Koster J, Rahmann S. Snakemake--a scalable bioinformatics workflow engine. Bioinformatics 2012;28(19):2520–2522.

41. Kivimäki M, Singh-Manoux A, Pentti J, et al. Physical inactivity, cardiometabolic disease, and risk of dementia: an individual-participant meta-analysis. Bmj Clin Res Ed 2019;365:1495.

42. Korologou-Linden R, Anderson EL, Howe LD, et al. The causes and consequences of Alzheimer’s disease: A Mendelian randomization analysis. Medrxiv 2019;2019.12.18.19013847.

43. Larsson SC, Traylor M, Malik R, et al. Modifiable pathways in Alzheimer’s disease: Mendelian randomisation analysis. Bmj 2017;359:j5375.

44. Østergaard SD, Mukherjee S, Sharp SJ, et al. Associations between Potentially Modifiable Risk Factors and Alzheimer Disease: A Mendelian Randomization Study. Plos Med 2015;12(6):e1001841. discussion e1001841.

45. Pierce BL, Burgess S. Efficient design for Mendelian randomization studies: subsample and 2-sample instrumental variable estimators. Am J Epidemiol 2013;178(7):1177–84.

46. Hemani G, Bowden J, Smith GD. Evaluating the potential role of pleiotropy in Mendelian randomization studies. Hum Mol Genet 2018;27(R2):R195–R208.

47. Gkatzionis A, Burgess S. Contextualizing selection bias in Mendelian randomization: how bad is it likely to be? Int J Epidemiol 2018;

